# Hyper-variability in Circulating Insulin and Physiological Outcomes in Male High Fat-fed *Ins1^-/-^:Ins2^+/-^* Mice in a Conventional Facility

**DOI:** 10.1101/031807

**Authors:** Arya E. Mehran, Nicole M. Templeman, Xioake Hu, James D. Johnson

## Abstract

Insulin is an ancient, multi-functional hormone with essential roles in glucose homeostasis and energy storage. Recently, our group has taken advantage of the ability to limit insulin secretion *in vivo* by reducing insulin gene dosage to demonstrate that insulin hypersecretion is a requirement for diet-induced obesity. Our previous studies employed male *Ins1*^+/−^:*Ins2*^−/−^ mice that exhibit a complete inhibition of diet-induced hyperinsulinemia relative to *Ins1*^+/+^:*Ins2*^−/−^ littermate controls, as well as female *Ins1^−/−^:Ins2^+/−^* mice with transient, partial reduction in circulating insulin relative to *Ins1^−/−^:Ins2^+/+^* littermates. In the present study, we sought to extend these studies to male *Ins1^−/−^:Ins2^+/−^* mice on the same chow and high fat diets. Surprisingly, while reduced *Ins2* gene dosage appeared capable of reducing *Ins2* mRNA, insulin protein levels in these mice were not significantly reduced. Moreover, there was a marked hyper-variability in circulating insulin levels within and between two independent cohorts of mice that persisted over at least the first year of life. In Cohort 1, we observed a paradoxical increase in body weight in some high fat-fed male *Ins1^−/−^*:*Ins2^+/−^* mice relative to *Ins1^−/−^:Ins2^+/+^* littermate controls. This phenomenon is consistent with the known satiety effects of insulin and our previous observations with *Ins2* can be expressed in the brain. Collectively, our data reveal unexpected complexity associated with the *Ins2* gene in male mice, and establish the *Ins2* gene as a candidate for studying the effects of modifier genes and/or environmental influences on gene-to-phenotype variability. Further studies are required to define the molecular mechanisms of this phenotypic hyper-variability and to define the role of reduced *Ins2* gene dosage in the brain.

## Introduction

Insulin genes are highly conserved, playing critical roles in glucose homeostasis in all species studied to date [1, 2]. Unlike humans, mice have two insulin genes, *Ins1* and *Ins2* [3]. Most studies have shown that *Ins1* is restricted to pancreatic β-cells, where it contributes to approximately 1/3 of the expressed and secreted insulin [3, 4]. The peptide product of the *Ins1* gene differs from that of the *Ins2* gene by two amino acids in the β-chain, at the B9 and B29 location, and is missing two amino acids in the connecting C-peptide [5]. *Ins1* also lacks an intron present in *Ins2* [5]. *Ins2* is the ancestral gene, with gene structure, parental imprinting, and a broad tissue distribution similar to human *INSULIN* [3, 4, 6]. Notably, there is evidence that both mouse *Ins2* and human *INSULIN* are expressed at low levels in within sub-populations of cells in the brain [4, 7]. The two murine insulin genes are partially redundant and capable of compensating for the loss of one another [8]. However, some studies, such as those comparing the effect of the expression of the *Ins1* versus *Ins2* in the thymus in the context of type 1 diabetes, have shown that the two genes are not entirely redundant [9]. Outside of type 1 diabetes (i.e. in conditions of relative normoglycemia), the effects of changed *Ins* gene dosage, and ultimately insulin levels, remain to be fully elucidated.

Studies of human populations and animal models of obesity have demonstrated that elevated levels of fasting insulin, known as hyperinsulinemia, precede weight gain [10-23]. Moreover, some studies have suggested that humans with class I allele VNTR in the *INSULIN* gene produce and release more insulin from the pancreatic islets and are also more susceptible to obesity [24-26], although this observation remains controversial [27]. On the other hand, studies of invertebrates with reduced insulin or insulin signalling have reported leaner, smaller bodies, along with increased lifespan [28, 29]. Similarly, studies in mammalian models, such as the Zucker fatty rats, have shown that treatment with diazoxide, a compound that reduces insulin secretion, results in reduced weight and improved glucose intolerance [30, 31]. Treatment of obese patients with diazoxide is also associated with weight loss in some small clinical trials [32, 33]. Lustig and his group found similar results using Octreotide, a somatostatin agonist that binds the sst5 somatostatin receptor, found on β-cells, which inhibits insulin release [34-36]. Therefore, such observations have raised the question of whether hyperinsulinemia itself is a primary defect in obesity. Recently, our group has extended the observations that a full complement of insulin genes appears to be required for substantial high fat diet-induced obesity in mammals [4, 37].

Here, we report on the phenotype of male *Ins1^−/−^:Ins2^+/−^* mice fed two different diets in a conventional facility. Surprisingly, the effect of *Ins2* gene dosage on circulating insulin peptide was highly variable in these mice, displaying strong cohort dependence. This precluded definitive conclusions about the effects of this gene on weight gain, but provide insight into the regulation and effects of insulin production from this locus.

## Materials and Methods

### Experimental Animals

The *Ins1^-/-^* and *Ins2^−/−^* mice were previously generated by Jacques Jami (INSERM) and are described elsewhere. [8] A neo cassette was used to disrupt the *Ins1* gene and replace most of its sequence. A β-geo (Neo/LacZ) cassette was used to disrupt most of the *Ins2* gene sequence. [8] We used DNeasy Blood and Tissue Kit (Qiagen, Valencia, CA) to isolate DNA from tail samples. The genotyping primers are detailed in Table 1.

**Table 1.**
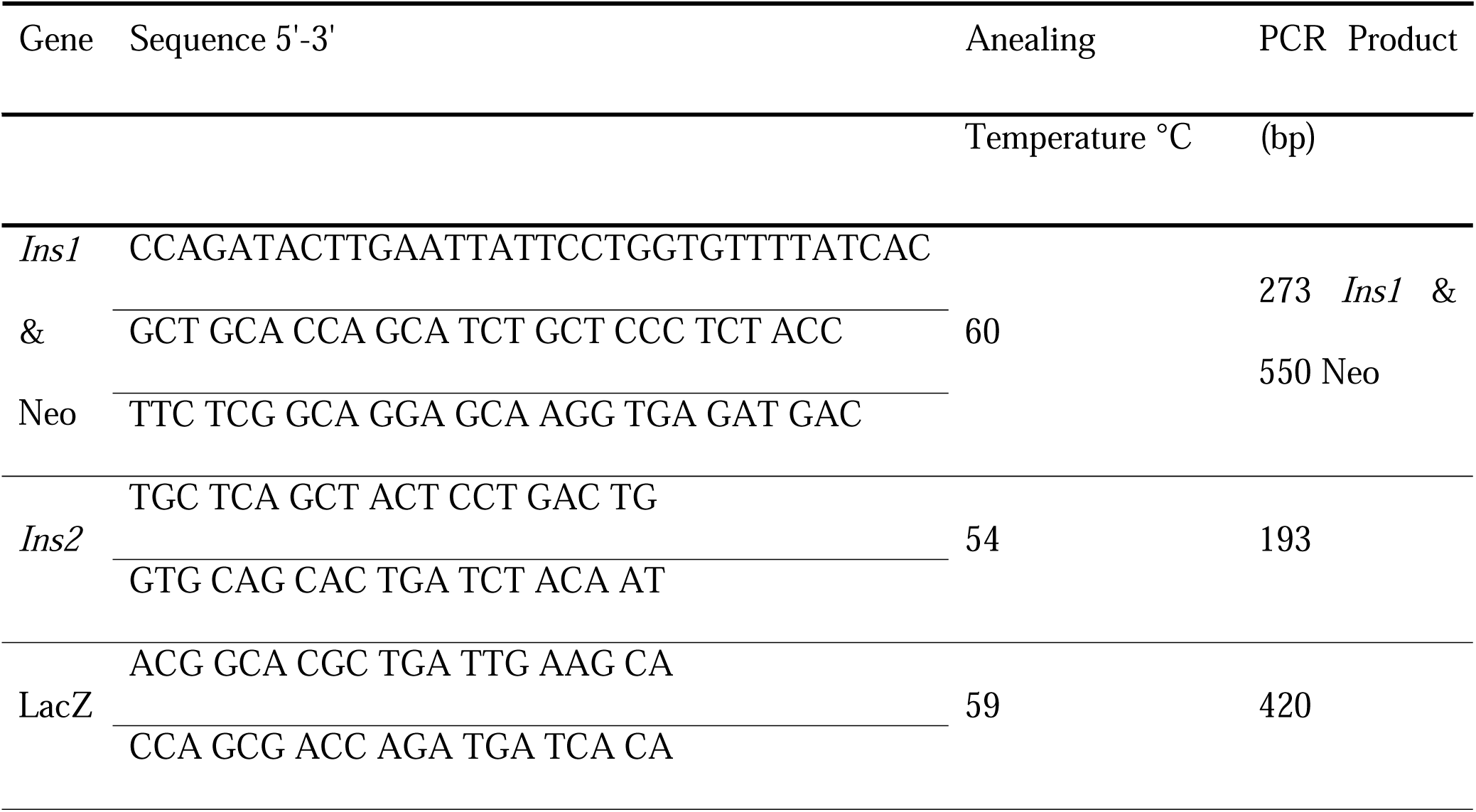
Primers used for genotyping the presence or absence of *Ins1* or *Ins2* alleles.

We divided the mice into two diet groups at weaning (3 weeks); one group was kept on the chow diet (CD; total calories = 3.81 kcal/g; 25.3% calories from fat, 19.8% calories from protein, 54.9% calories from carbohydrate; Catalog #5LJ5; PMI Nutrition International, St. Louis, MO) and we put the second group on a high fat diet (HFD; total calories = 5.56 kcal/g; 58.0% calories from fat, 16.4% calories from protein, 25.5% calories from carbohydrate; Catalog # D12330 Open Source Diets/Research Diets, New Brunswick, NJ). Diets are detailed in Table 2. One-year-old mice were scanned for whole body fat to lean mass ratio using NMR Spectroscopy at the 7T MRI Research Center at the University of British Columbia (Vancouver, BC).

**Table 2.** Comparison of the control medium fat and the high fat diets.

| Diet | 5015 (CD) | D12330 (HFD) |
| --- | --- | --- |
| Protein | 19.805 kCal% | 16.4 kCal% |
| Carbohydrate | 54.858 kCal% | 25.5 kCal% |
| Fat | 25.337 kCal% | 58.0 kCal% |
| Total Calories | 3.81 kCal/g | 5.55 kCal/g |
| Type of Fat | Lard | Hydrogenated Coconut Oil |

### Glucose Tolerance, Insulin Tolerance, and Hormone Secretion

We measured body weight and fasting glucose (OneTouch glucometer, LifeScan Canada, Burnaby, BC) weekly in four-hour fasted mice. The fasting was initiated at approximately 8 am (start of light cycle was at 7 am). For glucose tolerance tests, mice were injected intraperitoneally with 11.1 μL per gram of body weight of 18% glucose in 0.9% NaCl saline. For insulin tolerance tests, mice received 0.75 U of insulin (Lispro Humalog VL-7510 in 0.9% NaCl solution) per gram of body weight. Ultrasensitive mouse insulin ELISA kits (80-INSMSU-E01; ALPCO Diagnostics, Salem, NH) were used to measure serum insulin levels and leptin ELISA kits (90030; CrystalChem Inc., Downers Grove, IL) were used to measure serum leptin levels. Blood was collected from the tail vein.

### Metabolic Cage Analyses

We placed 8-week old mice (N = 3-5) from each group in PhenoMaster indirect calorimetry cages (TSE Systems Inc., Chesterfield MO) for three complete days. The cages also measured food, drink and body weight as well as activity using infrared beam grid in the x, y and z axes. All cages were contained in an environmental chamber to ensure constant temperature (21°C). The room’s light cycles were from 7 am - 7 pm. Data collected from the first 4 hours were not included in the study. The average of data collected from each of the 3 days were presented as a prototypical day for each genotype, as in our previous publications [4, 37].

### Tissue Collection and Analyses

At one year of age, mice were euthanized for the purpose of tissue collection. The following tissues were collected: pancreas, epididymal fat pads, soleus muscle, liver, brain, kidney, spleen, heart, thymus and tibia. Some samples were snap-frozen in liquid nitrogen and stored in −80°C freezer. The rest of the samples were fixed in 4% paraformaldehyde (PFA) for tissue sectioning. For removal of non-bone tissue the tibias were incubated in 2% KOH for for physical measurements. Sections were made serially at 5 μm thickness paraffin sections. The Child and Family Research Institute Histology Core Facility (Vancouver, BC) were responsible for making the sections. Pancreatic islets at 200 μm apart were stained with guinea pig anti-insulin and rabbit anti glucagon (Linco/Millipore). Using the insulin positive area morphology and hormone expression were approximated. The secondary antibodies of choice were Alexa Fluor 488 and 594 raised in goat (Life Technologies, Abtenau, Austria). The antibody dilutions were 1:100 for the primary antibodies and 1:400 for the secondary antibodies. Samples were incubated with primary antibodies overnight at 4 °C and one hour at room temperature with the secondary antibody. Vectashield solution with DAPI (Reactolab SA, Switzerland) was used as the mounting media. Imaging was done with a Zeiss 200M inverted microscope equipped with a 10x (1.45 numerical aperture) objective, individual filter cubes for each color, and a CoolSnap HQ2 Camera (Roper Scientific). Image analysis was done using the Slidebook software (Intelligent Imaging Innovations) as previously described [38].

### Statistical Analyses

Most results are expressed as means ± SEM. The area under the curve (AUC) was used to measure statistical significance in different groups for the glucose tolerance and insulin secretion tests and is described elsewhere [4]. The area over the curve (AOC) was used for insulin tolerance studies and are also described in detail elsewhere [4]. SPSS 15.0 software or Prism 5 (Graphpad) software was used to perform the statistical analyses. Two-way ANOVAs were used to compare factors of genotype and diet, or alternatively in the case of significant interactions, one-way ANOVAs with Bonferroni corrections were used. We used Levene’s test to validate homogeneity of variance. In all statistical analyses, if *p* < 0.05 the differences were considered to be significant.

## Results

### Effects of reducing *Ins2* gene dosage on *Ins2* mRNA, insulin production, and insulin secretion

We varied the *Ins2* gene dosage in mice lacking both alleles of *Ins1*, placing mice on a chow diet with moderate fat, or a diet high in fat (Fig. 1A). Deleting one of the two *Ins2* alleles reduced the *Ins2* mRNA levels in the *Ins1^−/−^:Ins2^+/−^* mice regardless of diet (Fig. 1B). Insulin content in isolated islets was not statistically different between genotypes, but showed a tendency to be decreased in *Ins1^−/−^:Ins2^+/−^* mice compared to littermate control *Ins1^−/−^:Ins2^+/+^* mice (*p* = 0.095; Fig. 1C). Immunofluorescent staining of islets from year-old mice (Fig. 1D) mirrored the measurements of insulin protein in isolated islets at 8 weeks (Fig. 1C). We did not detect significant differences in pancreatic β-cell area between groups (Fig. 1E), in contrast to *Ins1^+/−^:Ins2^−/−^* male mice in our previous report [4].

**Figure 1.**
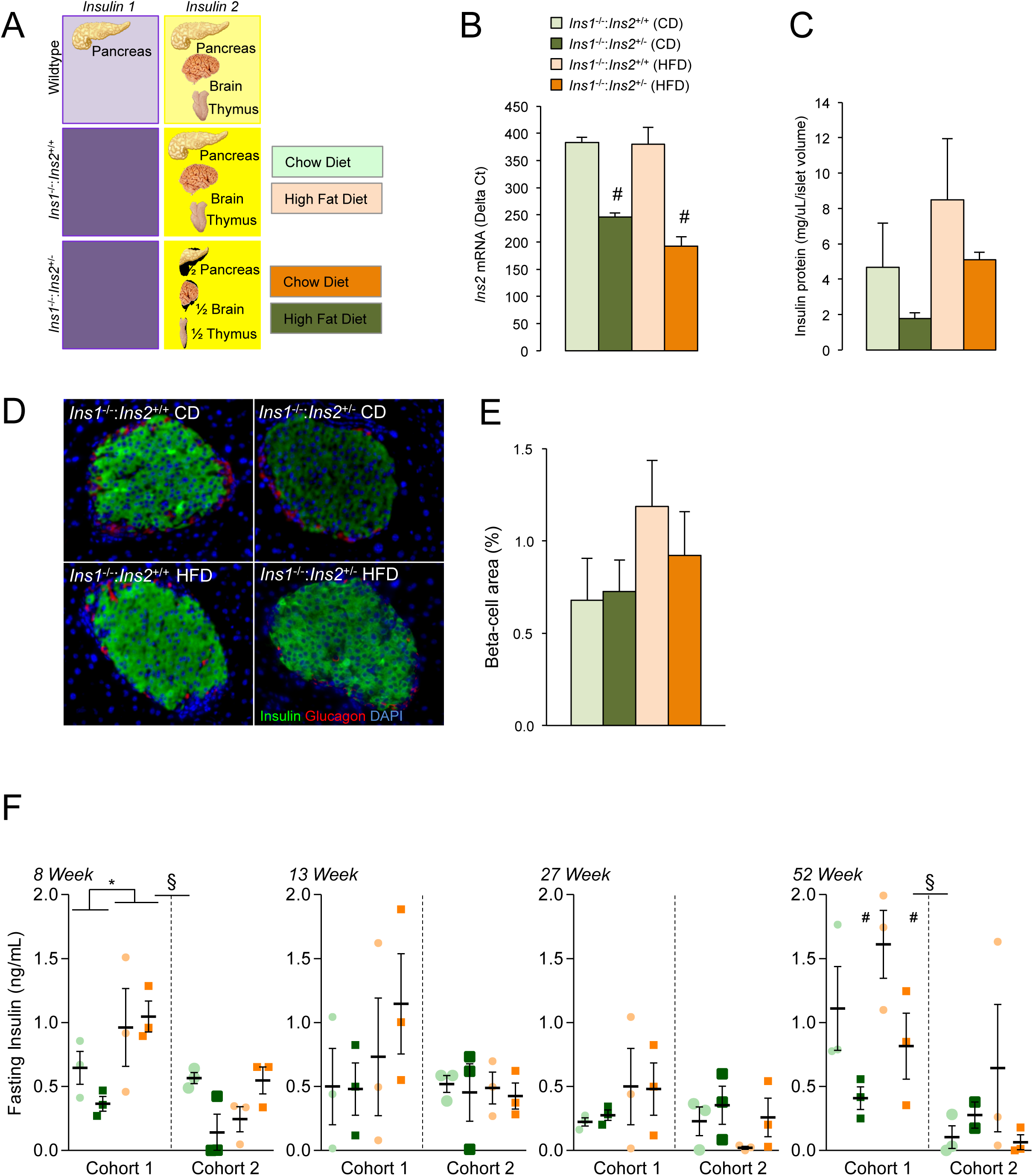
Reduced *Ins2* gene dosage and mRNA expression does not equate to consistently reduced insulin secretion in male *Ins1^−/−^:Ins2^+/−^* mice. **(A)** Experimental design for mice with varying *Ins2* gene dosage on an *Ins1* null background. **(B)** Proportionally reduced *Ins2* mRNA in *Ins1^−/−^:Ins2^+/−^* mice regardless of diet in islets isolated from 8 week old mice (n = 3-4 per group). As expected, *Ins1* mRNA was not found in these samples (not shown). **(C)** Insulin protein content in 30 size-matched islets isolated from 8 week old mice (n = 3-4 per group). The samples in Panels A-C, which required euthanasia for collection, were from neither Cohort 1 nor Cohort 2, which were followed for 1 year. **(D,E)** Insulin immunoreactivity in pancreas secretions from cohort 1 mice collected at 1 year of age and used to assess beta-cell area. **(F)** Circulating insulin was measured at 4 time points in both Cohort 1 (n = 3) and Cohort 2 (n = 2-3). *p ≤* 0.05 denoted by * for CD vs HFD, # for *Ins1^−/−^:Ins2^+/+^* vs *Ins1^−/−^:Ins2^+/−^*, and § for cohort 1 vs cohort 2.

We next measured fasting insulin, which is the product of the number of β-cells and their basal insulin exocytosis. Interestingly, fasting insulin was hyper-variable in these mice, with only cohort 1 exhibiting significant HFD-induced elevated insulin at 8 weeks of age, and lowered fasting insulin in year-old *Ins1^−/−^:Ins2^+/−^* mice compared to *Ins1^−/−^:Ins2^+/+^* mice (Fig. 1F). Fasting insulin was also significantly different between cohorts at 8 and 52 weeks of age (Fig. 1F). Collectively, these data demonstrate that while reducing *Ins2* gene dosage has the expected effect of reducing *Ins2* mRNA, compensatory post-transcriptional mechanisms appear to have resulted in wide variation in circulating insulin levels in these mice.

### Glucose Homeostasis in *Ins1^−/−^:Ins2^+/−^* Mice

Glucose homeostasis was tracked over 1 year in *Ins1^−/−^:Ins1^+/−^* mice and *Ins1^−/−^:Ins1^+/+^* littermate controls. No significant differences in fasting blood glucose levels between groups were observed throughout the experiment (Fig. 2A; both cohorts pooled). Consistent with the observed fasting insulin results (Fig. 1F), there were significant differences between cohorts for glucose-stimulated insulin secretion values at 7 and 52 weeks of age. However, in spite of differences in value magnitude, there tended to be similar patterns of the effects of diet and genotype across both cohorts for stimulated insulin secretion and blood glucose response to intraperitoneal glucose or insulin, and so the cohorts were pooled for these data. Glucose-stimulated insulin secretion was higher in high fat-fed mice compared to chow-fed mice at 7 weeks of age, but there were no statistically significant differences in stimulated insulin secretion detected between *Ins1^−/−^:Ins2^+/−^* mice and their *Ins1^−/−^:Ins2^+/+^* littermate controls at any time point (Fig. 2B). This indicates that a single allele of the *Ins2* gene is sufficient to generate enough insulin to mount an appropriate response to glucose in these male mice, unlike the previously described male *Ins1^+/−^:Ins2^−/−^* mice [4] or female *Ins1^−/−^:Ins2^+/−^* mice [37]. Effects of genotype on glucose tolerance were very modest, with *Ins1^−/−^:Ins2^+/−^* mice exhibiting slightly worsened glucose tolerance than their *Ins^−/−^:Ins2^+/+^* littermates only at 25 weeks (Fig. 2C). In addition, a minimal degree of HFD-induced glucose intolerance was observed by 52 weeks of age (Fig. 2C). Interestingly, high fat-fed *Ins1^−/−^* mice (regardless of *Ins2* gene dose) were paradoxically insulin hypersensitive at all of the time points from 12 weeks onward (Fig. 2D). On chow diet, 6 week-old mice with reduced *Ins2* gene dosage tended to be hypersensitive to exogenous insulin, but this was not observed in older mice (Fig. 2D).

**Figure 2.**
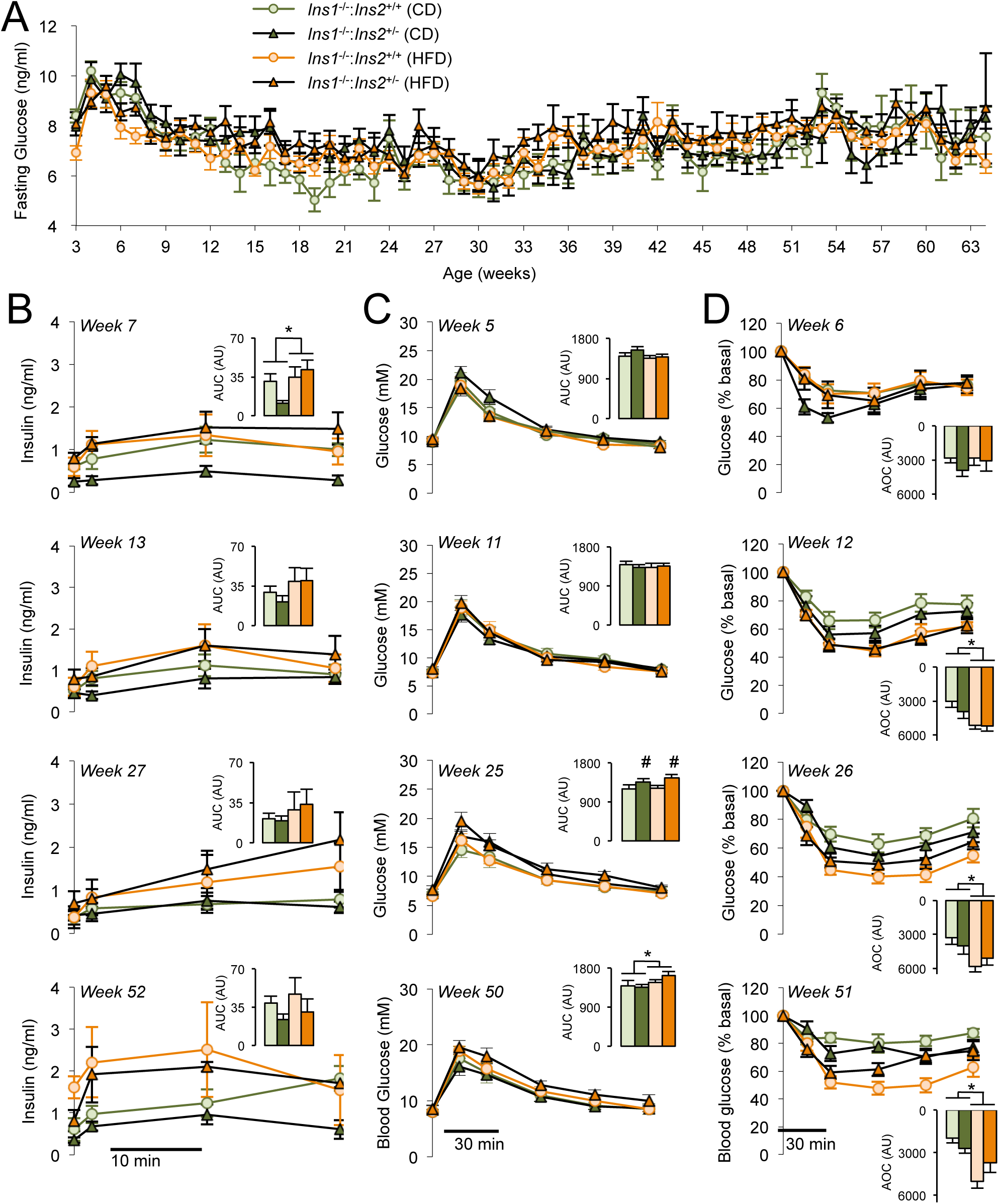
Insulin secretion and glucose homeostasis in high fat fed *Ins1^−/−^:Ins2^+/−^* mice. **(A)** Weekly blood glucose after a 4-hour fast. **(B)** Glucose-stimulated insulin release. Insets show area under the curve (AUC). **(C)** Intraperitoneal glucose tolerance. Insets show area under the curve (AUC). **(D)** Insulin tolerance (0.75 U/g) after four hours of fasting. Insets show area over the curve (AOC). n = 6-14 per group. Cohort data are pooled. *p ≤* 0.05 denoted by * for CD vs HFD and # for *Ins1^−/−^:Ins2^+/+^* vs *Ins1^−/−^Ins2^+/−^*.

### Cohort-dependent Effects of *Ins2* Gene Dosage on Diet-induced Obesity

Comparing the effects of a chow diet and high fat feeding on *Ins1^−/−^:Ins2^+/−^* mice and control *Ins1^−/−^:Ins2^+/+^* mice revealed a complex, cohort-dependent gene-environment interaction. In the first cohort, *Ins1^−/−^:Ins2^+/+^* mice did not gain weight on a high fat diet (Fig. 3A), leading us to speculate that the hypersecretion of the *Ins1* gene product could have been required for the proper storage of lipids in adipose tissue. Remarkably, however, the majority of *Ins1^−/−^:Ins2^+/−^* mice in the first cohort exhibited striking weight gain on the high fat diet (Fig. 3A). Surprisingly, these somewhat paradoxical differences were not observed in a second cohort (Fig. 3B). Together, these data illustrate a hyper-variability of the physiological response to reduced *Ins2* gene dosage in these male mice.

**Figure 3.**
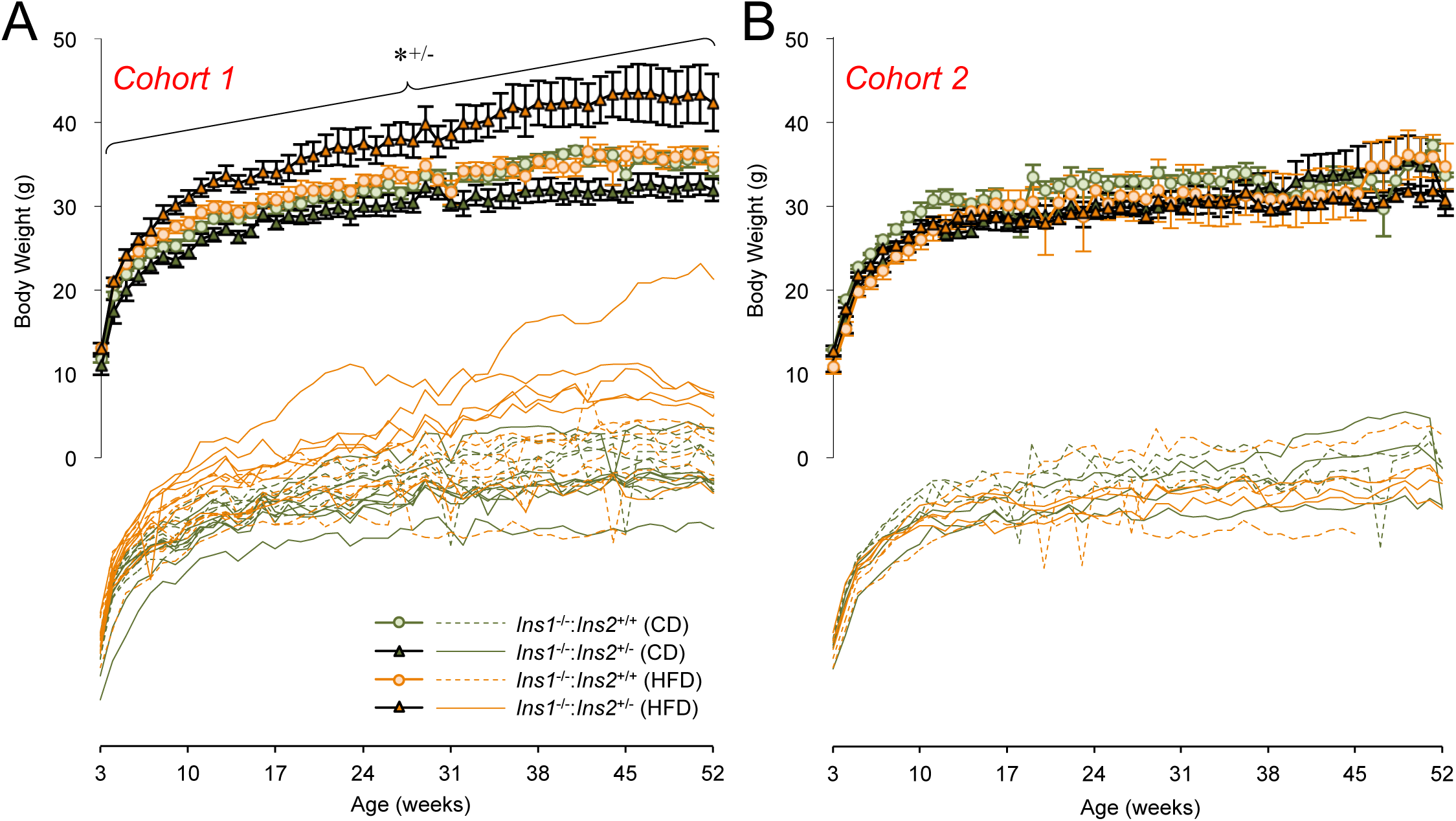
Cohort-dependent and diet-dependent effects on body weight in *Ins1^−/−^:Ins2^+/−^* mice on high fat diet. Pooled body weight tracked weekly for 1 year in Cohort 1 **(A)** and Cohort 2 **(B)**. Individual body weight are also shown across 1 year in Cohort 1 **(A)** and Cohort 2 **(B)**. In cohort 1, n = 6-8, and in cohort 2, n = 3. \**^+/−^* denotes *p* < 0.05 for CD-vs HFD-fed *Ins1^−/−^*:*Ins2^+/−^* mice.

### Characterization of Weight Gain in the First Cohort of *Ins1^−/−^: Ins2^+/−^* Mice

Weight gain in the first cohort of high fat-fed *Ins1^−/−^:Ins2^+/−^* mice, relative to the other groups, appears to have been due to a combination of increased adiposity and increased somatic growth of various organs (Figs. 4A-F). An unusual heterogeneity in adipocyte size was observed in all groups of *Ins1^−/−^* mice (Fig. 4A), reminiscent of mice lacking adipocyte insulin receptors [39, 40]. Circulating leptin tended to be proportional to epididymal fat pad weight and fat-to-lean ratio (Figs. 4 B,C,F). We did not detect differences in the circulating free fatty acids between groups in this cohort (Fig. 4D). Together, these data suggest that, in some currently un-defined conditions, reducing *Ins2* expression can be associated with facilitating greater weight gain, via a mechanism that is active specifically in the context of a high fat diet.

**Figure 4.**
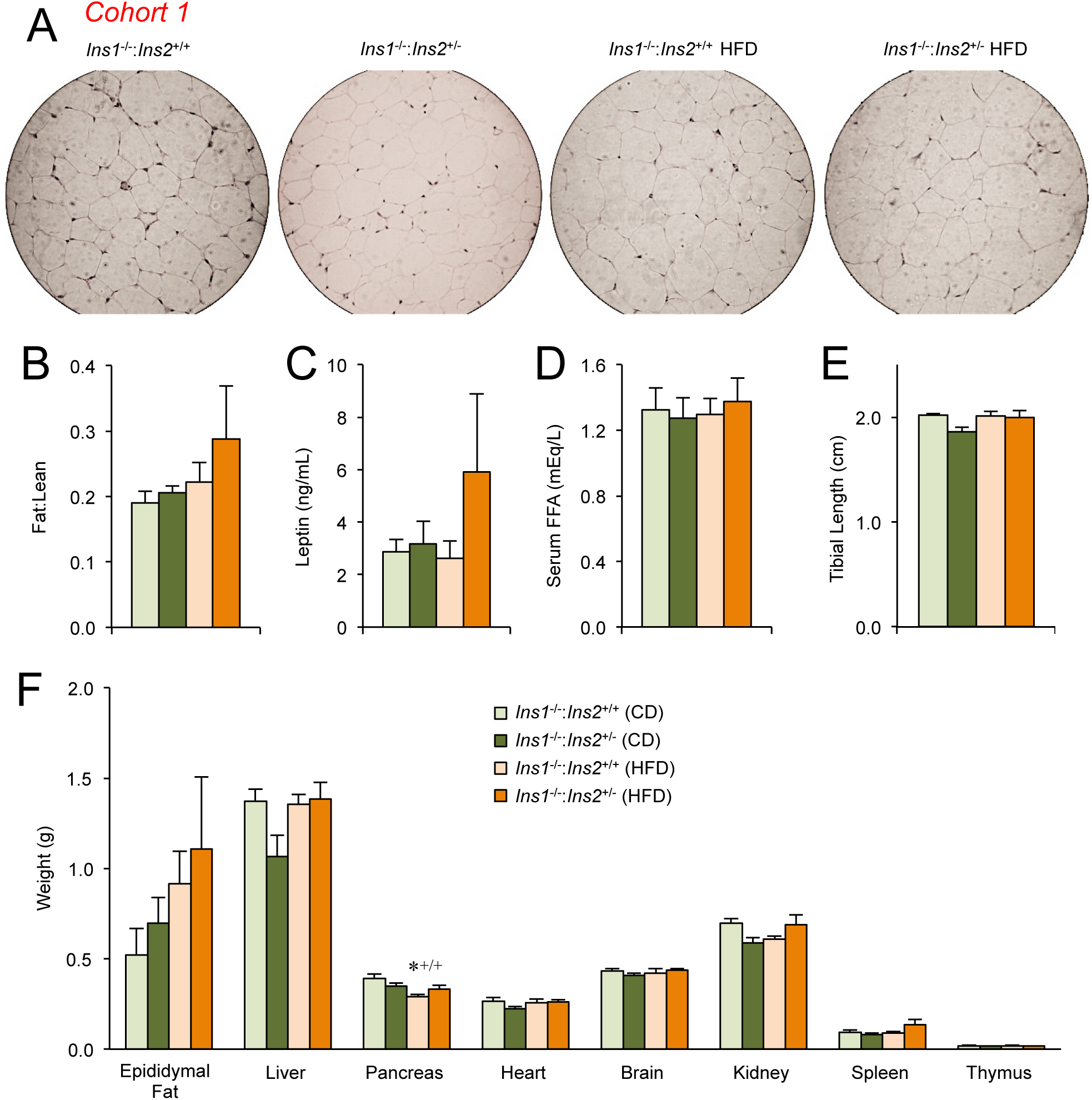
High Fat Diet-dependent Adiposity in *Ins1^−/−^:Ins2^+/−^* Mice From Cohort 1. **(A)** Hematoxylin and eosin-stained epididymal fat revealed heterogeneity in adipocyte size (representative image from 1 of 3 mice assessed per group). **(B)** Whole body fat to lean ratio measure with NMR (n = 3). **(C)** Circulating leptin levels (n = 3). **(D)** Serum free fatty acids (n = 3). **(E,F)** Tibial length and tissues weights from 1 year old mice (n = 6-8). *^+/+^ denotes *p* < 0.05 for CD-vs HFD-fed *Ins1^−/−^*:*Ins2^+/+^* mice. Data are collected from Cohort 1.

Given the statistical similarity between genotypes for the average circulating insulin levels prior to one year of age, these observations hint at the possibility of altered local effects of *Ins2* in the brain. It is well established that insulin can act in the brain as a satiety factor [41, 42]. We have confirmed that *Ins2* is expressed in multiple regions of the brain that can potentially control and project to feeding, reward and memory centers, raising the possibility that central *Ins2* gene expression may regulate food intake [4]. In this first cohort, where reduced *Ins2* gene dosage was associated with increased weight gain on high fat diet, there was a tendency for *Ins1^−/−^:Ins2^+/−^* mice to show elevated food intake on HFD versus CD, whereas lean *Ins1^−/−^:Ins2^+/+^* littermates tended to to reduce caloric intake in response to high fat feeding (Fig. 5A). Other parameters such as activity and energy expenditure were not statistically different between any of the groups (Figs. 5B-E). Taken together, these experiments indicate that, within this cohort, reduced *Ins2* gene dosage led greater weight gain on high fat diet, which may have been associated with increased relative food intake.

**Figure 5.**
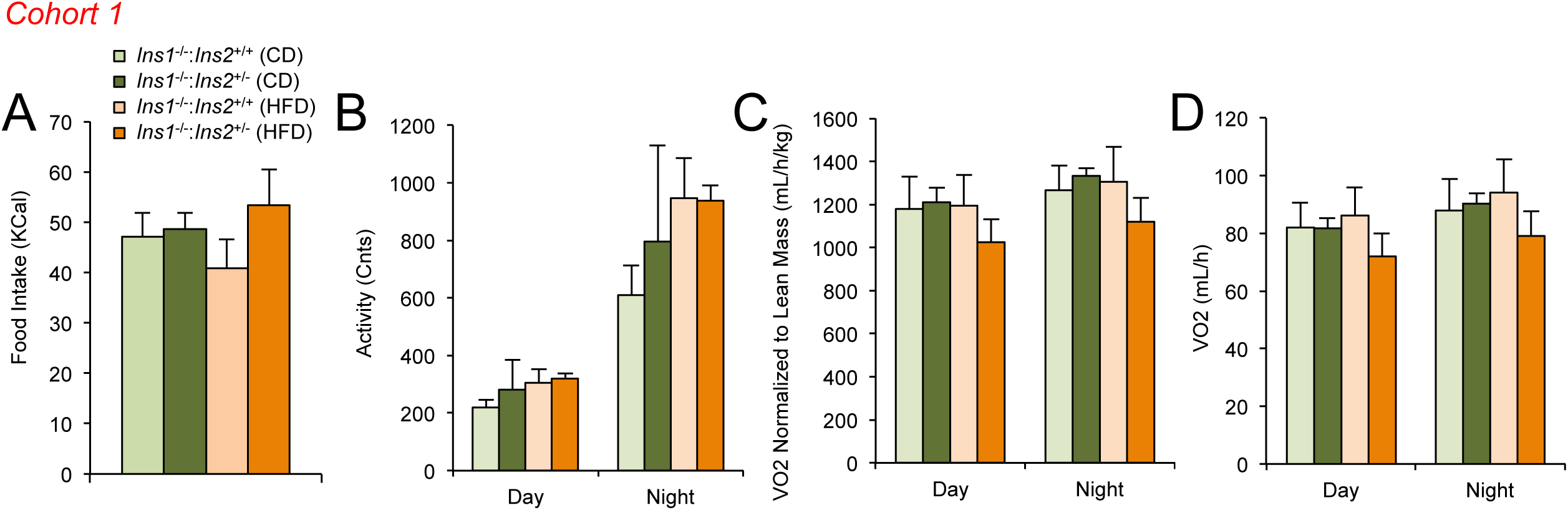
Food intake and metabolic parameters in Cohort 1. **(A-D)** Food intake, activity, and oxygen consumption (normalized to body weight, or total oxygen consumption) were measured over 3 days in indirect calorimetry cages at 8 weeks of age in male mice from Cohort 1 (n = 5-14).

## Discussion

The present study was initiated to investigate the effects of reduced *Ins2* gene dosage, in the absence of *Ins1*, on glucose homeostasis and weight gain in male mice in the context of chow and high fat diets. To our surprise, reduced *Ins2* gene dosage did not translate into consistent differences in circulating insulin. We observed a large degree of variability within and between two cohorts of mice. These observations are in contrast to our experience with female mice of the same genotypes on the same diets, and in contrast with our experience reducing *Ins1* gene dosage in male mice [4, 37].

Insulin is the most studied hormone in biology, yet the present study provides new insight into the range of physiological functions that can be modulated by insulin. Specifically, our data point to somewhat unique roles for, and post-transcriptional regulation of, the *Ins2* gene, relative to the *Ins1* gene. Elegant studies in flies and worms have demonstrated that deleting specific insulin genes increases lifespan and prevents diseases associated with adiposity, and clearly indicate that individual insulin-like peptide genes have distinct physiological functions despite signalling through a single receptor [28, 29]. It has been previously shown that the mouse *Ins1* and *Ins2* genes have opposing effects on type 1 diabetes incidence in the NOD mouse due to the induction of thymic tolerance by *Ins2*, [43, 44]. Specifically, *Ins2* expression was associated with reduced incidences of type 1 diabetes and the reverse was true for the expression of the *Ins1* gene [43, 44]. Our previous observation that the pancreatic-specific *Ins1* is dose-dependently required for diet-induced obesity in male mice [4], defines the first specific role for *Ins1* outside the context of type 1 diabetes. Similarly, a follow-up study revealed that a modest and transient reduction in circulating insulin in female *Ins1^−/−^: Ins2^+/−^* mice relative to *Ins1^−/−^:Ins2^+/+^* littermate controls was sufficient to provide long-term protection from diet-induced obesity [37]. In the present study, the high fat diet was unable to consistently increase fasting insulin or beta-cell mass in a statistically significant manner. This meant that we were unable to formally test the hypothesis that reducing hyperinsulinemia by reducing *Ins2* gene dosage might protect these mice from high-fat diet-induced obesity, as we could in our previously published studies [4, 37]. Clearly, additional studies with greater statistical power and more diet groups would be required to formally rule in or out a role for hyperinsulinemia stemming from the *Ins2* gene in diet-induced obesity.

One interesting observation from the present study was that male *Ins1^+/−^:Ins2^−/−^* from the first cohort were heavier than their *Ins1^+/−^:Ins2^−/−^* littermates fed the same high-fat diet. Considerable mouse-to-mouse heterogeneity was observed in these measurements, but they were relatively consistent within each mouse over time. Taken at face value, these observations suggest the possibility that *Ins2* expression in the brain made have played a role in this weight gain, because only the *Ins2* gene is robustly expressed in the brain [4], and because we only detected modest and late-onset differences between genotypes in circulating insulin levels. The central nervous system is known to play important roles in peripheral energy homeostasis and body weight regulation [45-47]. For example, it has been reported that insulin receptor knockout in the brain leads to increased high fat food intake and obesity [48], suggesting the possible presence of a local signalling network. The presence of small amounts of insulin protein and mRNA in the mammalian brain has long been reported [49](reviewed in [4]). The production of *Ins2* mRNA and protein in specific brain regions was confirmed by our group using *Ins1^−/−^* and *Ins2^−/−^* as negative controls, and *Ins2^βGal/+^* mice as positive staining controls [4].

Since insulin has been proposed to be a satiety factor [50], downregulation of insulin expression and/or action may be expected to increase food intake. Consistent with this notion, *Ins1^−/−^:Ins2^+/−^* mice in our first cohort tended to show increased high fat food intake and were obese when compared to the high fat-fed *Ins1^−/−^:Ins2^+/+^* littermate controls. Although our experimental treatment was expected to reduce *Ins2* gene dosage in multiple tissues, including the pancreas, thymus and brain, several lines of evidence could potentially hint that a partial reduction of brain *Ins2* was associated with the diet-dependent hyperphagia and obesity, at least in some specific conditions. Other investigators have shown that *Ins2* knockout in the thymus had no effect on body weight [51], further suggesting that the brain could have play a role in this phenomenon. Our work should open new avenues for investigating the biology of insulin in neurons and their connections. However, it is imperative that these data are interpreted with caution, as the second cohort of *Ins1^−/−^:Ins2^+/−^* mice did not gain additional weight with high fat feeding. A complete understanding of the role of *Ins2* in the central nervous system will require conditional *Ins2* knockout mice.

The phenotypes we observed in the present study were hyper-variable, sex-specific, and diet-dependent with respect to reduction of the *Ins2* gene. The animal facility where this work was done no longer exists, so it is impossible to formally repeat these studies in the same environment. Similar studies were also conducted in a modern specific-pathogen free facility, and cohort-dependent variability is described in a companion paper. Remarkably, in this distinct animal facility we observed opposite outcomes for the effect of *Ins2* gene dosage on weight gain than those which were shown in the current experiments. Collectively, the experience of our laboratory is that modulation of the *Ins2* gene, in the absence of *Ins1*, leads to striking variability in body weight. The source of this variability will be investigated by our collaborators specializing in epigenetics and translational control. Phenotypic hyper-variability is a poorly understood phenomenon in biology, but our studies illustrate that varying *Ins2* gene dosage could be a useful model with clear differences in physiological outcomes.

## Acknowledgements

We thank Professor Jami for donating the mouse strains.

